# Targeting effector pathways in RAC1^P29S^-driven malignant melanoma

**DOI:** 10.1101/750489

**Authors:** Cristina Uribe-Alvarez, Sandra Lucía Guerrero-Rodríguez, Jennifer Rhodes, Alexa Cannon, Jonathan Chernoff, Daniela Araiza-Olivera

## Abstract

Malignant melanoma is characterized by mutations in a number of driver genes, most notably *BRAF* and *NRAS*. Recently, genomic analyses revealed that 4-9% of sun-exposed melanoma bear activating mutations in *RAC1*, which encodes a small GTPase that is known to play key roles in cell proliferation, survival, and migration. The RAC1 protein activates several effector pathways, including Group A p21-activated kinases (PAKs), phosphoinositol-3-kinases (PI3Ks), in particular the beta isoform, and the serum-response factor/myocardin-related transcription factor (SRF/MRTF). Having previously shown that inhibition of Group A PAKs impedes oncogenic signaling from RAC1^P29S^, we here extend this analysis to examine the roles of PI3Ks and SRF/MITF in melanocytes and/or in a zebrafish model. We demonstrate that a selective Group A PAK inhibitor (Frax-1036) and certain PI3Ks inhibitors (BKM120, TGX221, GSK2636771) impede the growth of melanoma cells driven by mutant RAC1 but not mutant BRAF, however other PI3K inhibitors, including PI3Kα-selective inhibitors are less effective. Similar results were seen *in vivo*, using embryonic zebrafish development as a readout, but now including an SRF/MRTF inhibitor (CCG-203971). These results suggest that targeting Group A PAKs and/or SRF/MRTF represent promising approach to suppress RAC1 signaling in malignant melanoma.

## Introduction

Malignant melanoma is a highly aggressive cancer associated with poor overall survival. Recent genomic analyses have uncovered a variety of new driver mutations in malignant melanoma including an activating mutation in *RAC1*.^1, 2^ *RAC1* encodes a small ubiquitously expressed GTPase known to play key roles in embryonic development, immune response, cell proliferation, survival, and rearrangement of cytoskeleton by actin filament remodeling.^3–5^ In sun-exposed cutaneous melanomas, *RAC1^P29S^* is the third most common oncogenic driver mutation, following *BRAF^V600E^* and *NRAS^Q61L/K/R^*.^1, 2^ Analysis of the RAC1^P29S^ structure and biochemical studies have shown that the proline to serine substitution in the hydrophobic pocket of the switch I domain of the GTPase results in an increased cycling rate from the GDP-bound inactive state to the GTP-bound active state, triggering downstream effectors and promoting melanocyte proliferation and migration^1^.

While RAC1 itself represents a challenging therapeutic target, its effectors might be more tractable. Many of RAC1’s downstream biological effects are propagated by p21-activated kinases (PAKs). PAKs are serine/threonine specific intracellular kinases that phosphorylate downstream effector substrates regulating apoptosis, cell motility, cell morphology and cytoskeleton rearrangement.^6^ Overexpressed PAKs result in oncogenic effects including increased cell proliferation, cell cycle progression, evasion of apoptosis, angiogenesis, and promotion of invasion and metastasis.^7^ RAC1^P29S^ induces PAK1 activation, which in turn phosphorylates and activates MEK1 at the Serine 298 site, facilitating ERK activation and transcription of various target genes. The PAK/MEK/ERK pathway is essential for RAS-driven transformation in a mouse model of skin cancer^8^ and represents a potential therapeutic target for sun-driven melanomas. Zebrafish embryos injected with *RAC1^P29S^* mRNA displayed abnormal development and PAK and ERK elevated activity^9^. Defective growth was reversed by PAK and MEK inhibitors, suggesting that these may be useful to prevent the developmental effects of RAC1^P29S^ mutations^9^. In addition, we and others have reported that tumors and human melanoma cell lines bearing *RAC1^P29S^* mutations are resistant to BRAF inhibitors but are sensitive to PAK and MEK inhibitors.^9, 10^

In addition to PAKs, PI3Ks represent a second recognized effector for RAC1^11^. PI3Ks are lipid signaling kinases that play key regulatory roles in cell survival, proliferation and differentiation.^12^ Class I PI3Ks can be divided into two families based on their regulation mode: Class IA are heterodimeric proteins composed by a p110 catalytic subunit (PI3Kα/p110α, PI3Kβ/p110β or PI3Kδ/p110δ) whose enzymatic activity depends on their binding to regulatory p85 subunit (p85α, p55α, p50α, p85β or p55γ); and Class IB which do not need to interact with a regulatory subunit to be active (PI3Kγ/p110γ)^12,13^. ‘ PI3Ks transduce external signals from growth factors and cytokines into phospholipids that activate various downstream effector pathways such as the serine/threonine kinase AKT, and guanine-nucleotide exchange factors (GEFs).^14^

The PI3K/AKT pathway is an important regulator of normal cell physiology and is observed to be activated in 70% of sporadic melanomas including those containing the *RAC1^P29S^* mutation.^15–18^ Activated AKT phosphorylates its protein targets inhibiting apoptosis and promoting cell survival. The AKT protein kinase family consists of three isoforms, while the targeted inhibition of AKT1 and AKT2 show little effect on combating melanoma, AKT3 inhibition resulted in increased apoptosis, reduced cell survival and decreased tumor development providing a new therapeutic target for patients with advanced stages of melanoma.^15, 17^

Recent transcriptome analysis from RAC1^P29S^ melanocytes identified an enrichment of the SRF transcription factor targets and the epithelial to mesenchymal transition genes (EMT).^11^ The nuclear transcription factor serum response factor (SRF) and its myocardin-related transcription factor (MRTF) co-factor are regulated by actin polymerization. Upon GTPase activation of the WAVE regulatory complex (WRC), the ARP2/3 complex catalyzes the polymerization of G-actin into F-actin liberating the SRT/MRTF transcription factor which may now bind to DNA and induce the transcription of epithelial to mesenchymal transition (EMT) genes. SRF/MRTF inhibitors could therefore represent an attractive approach to tackling resistance in melanoma bearing the *RAC1^P29S^* mutation.^11^

Vemurafenib and dabrafenib, a BRAF and a MEK inhibitor, respectively, inhibit growth and promote tumor regression in BRAF^V600E^ mutant melanomas.^19–21^ The effect of these drugs is limited due to an eventual reactivation of the MAPK pathway and the PI3K-AKT signaling pathway that lead to drug resistance.^22–28^ BRAFi and AKTi combinatorial therapies in human melanoma cells display promising effects on abatement of tumor growth. Additionally, combination of MEK, BRAF and AKT inhibitors delayed signs of drug resistance.^29^ In melanoma cells, the combination of PI3Kβ and PI3Kα inhibitors is necessary to inhibit the PI3K signaling in long-term treatments.^30^ BKM120, another PI3K inhibitor, prevented AKT activation, cell cycle arrest in the G2-M phase and induced apoptosis in human melanoma cells that metastasize to the brain in *in vitro* and *in vivo* assays. The combination of BKM120 with the MEK inhibitor binimetinib, further inhibited melanoma cell line proliferation.^31^ PAK and MEK inhibitors may also cause tumor regression, loss of ERK and AKT activity in a RAS-mediated skin cancer mice model^8^.

Here, we compare the results of inhibiting three distinct classes of RAC1 effectors – PAKs, PI3Ks, and SRF/MRTF – on the growth and survival of RAC1-mutant melanoma cell lines and on RAC1-driven changes in zebrafish embryonic development.

## Materials and Methods

### Reagents

PI3K inhibitors: TGX221, GSK2636771, AS252424, GSK2269557, BKM120 and BYL719; AKT 1/2/3 inhibitor: MK 2206; and MRTF/SRF inhibitor: CCG-203971 were purchased from Selleckchem. PAK1 inhibitor Frax-1036 was generously provided by Genentech.

### Cell culture

501mel, YUROB, YUFIC, YURIF and YUHEF melanoma cell lines were generously provided by Ruth Halaban (Yale University) and maintained in OptiMEM media (Invitrogen, Carlsbad, CA, USA) supplemented with 5% FBS and penicillin/streptomycin. 451Lu, WM1791 and WM1960 were generously provided by Meenhard Herlyn (Wistar Institute) and maintained in 80% MCDB153, 20% Leibovitz’s L-15, supplemented with 2% FBS, 5 μg/ml insulin, and 1.68 mM CaCl_2_. All cells were cultured in a humidified incubator with 5% CO2 at 37°C. Cells were tested for mycoplasma and authenticated by sequencing the *BRAF, PREX2*, and *RAC1* genes.

### Cell viability (mitochondrial activity)

Melanoma cell lines were plated in 96-well plates at 5000 cells/well in the corresponding medium. After 24 hours, the cells were treated with increasing concentrations of PI3K inhibitors. Cell viability was evaluated after 72-hour incubation with drugs. Culture media was replaced by 100 μL of MTT (3-(4,5-dimethylthiazol-2-yl)-2,5-diphenyltetrazolium bromide) to a final concentration of 0.33 mg/mL dissolved in culture media and incubated at 37 °C with 5% CO_2_ for 1 h in the dark. Supernatant was removed and replaced with DMSO to dissolve formazan crystals. Absorbance value was measured at 570 nm. Experiments were done in triplicate and 0. 1% DMSO was used as negative control.

### Immunoblotting

Western blot assays were performed using standard techniques. Primary antibodies used in this study were: antiphospho-PAK1 (pSer144) (#2606), -phospho-ERK1/2 (pThr202/pTyr204) (#9101) and -phospho-AKT (Ser473) from Cell Signaling Technology.

### Proliferation Assays

Proliferation was evaluated with the xCELLigence technology (Acea Bioscience, San Diego, CA, USA, distributed by Roche) in E-16-well plates. Melanoma cells were monitored during 72 h. The impedance value of each well was automatically monitored by microelectrodes placed on the bottom of plates. The impedance changes detected were proportional to the number of adhering cells and expressed as the cell index value. The experiments were conducted in triplicate.

### Wound healing assay

Melanoma cells were seeded on six-well plates and manually scraped with a 200-μL pipette tip. The cells were washed once with growth media, and then grown in fresh growth media with the IC50 of the inhibitors for 24 h. Images were acquired at 100x magnification using an EVOS fluorescence microscope and the number of cells that cross into the wound area from their reference point at time zero was analyzed. To determine the migration rate of the cells, the wound areas were quantified using ImageJ software.

### Zebrafish Microinjection Experiments

Wild-type AB zebrafish embryos were collected and maintained in the Zebrafish Core Facility of Fox Chase Cancer Center under standard conditions^39^. Capped mRNA was obtained as follows: RAC1 (P29S) cDNA was subcloned into the pSGH2 vector. Transcripts were made *in vitro* for antisense *(Hind*III and T3 RNA polymerase) or sense (*Sac*II and SP6 RNA polymerase) full-length mRNA using the mMessage Machine kit (Ambion). mRNA was injected directly in the cytoplasm of one-cell-stage embryos using a nitrogen-powered Picospritzer III injector (Intracel) conjugated to a Nikon SMZ 1000 stereomicroscope at a final concentration of 35 ng/μl, as described previously.^40^ 4 hpf mRNA injected embryos were incubated in E3 medium containing 1 μM of PI3K, PAK1, AKT or Rho/MRTF/SRF inhibitors for 1 h and then washed thoroughly. Control embryos were incubated in E3 with DMSO at the same final concentration as the small molecule inhibitors. To analyze zebrafish morphology, 24 h dechorionated embryos were placed on a glass depression slide in 1% methylcellulose to stabilize the embryo. Morphology was assessed visually using a light transmission Nikon SMZ 1500, and representative images were recorded using a Nikon digital sight DS Fi1 camera.

All animal procedures were performed in accordance with IACUC guides and regulations.

### Statistics

Statistical analyses were carried out using the paired Student’s t-test. All values reflect the mean ± S.E.M, with a significance cutoff of P<0.05. All statistical analyses were completed in GraphPad Prism 6.0 or 7.0 (La Jolla, CA, USA).

## Results

### Cytotoxic effect of PI3K inhibitors on Melanoma cell lines

Hotspot mutations of *BRAF, NRAS* and *RAC1* genes have been identified as the most frequent drivers in melanoma. In this study we analyze the effect of diverse inhibitors on melanoma cell lines with different mutations. Table 1 shows the target of each small molecule inhibitor.

**Table 1.** PI3K selective inhibitors and biological activity.

| Inhibitor | Target |
| --- | --- |
| BKM120<br>(Buparlisib) | pan-PI3K<sup>32</sup> |
| BYL719<br>(Alpelisib) | PI3Kα, minimal effect on PI3K$\beta/\gamma/\delta$<sup>33, 34</sup> |
| TGX221 | PI3Kβ, minimal effect on PI3Kα<sup>35</sup> |
| GSK2636771 | PI3Kβ<sup>36, 37</sup> |
| AS-252424 | PI3Kγ, minimal effect on PI3K$\delta/\beta$<sup>38</sup> |
| GSK2269557<br>(Nemiralisib) | PI3Kδ<sup>39</sup> |
| MK-2206 2HCl | AKT1/2/3<sup>40, 41</sup> |
| CCG-203971 | MRTF/SRF<sup>42</sup> |

Viability, proliferation, migration and signaling assays were performed in 8 different cell lines: YUROB (WT), 501mel *(BRAF^V600E^*), YURIF *(BRAF^V600K^/RAC1^P29S^)*, 451Lu *(BRAF^V600E^)*, YUFIC *(N-RAS^Q61R^)*, YUHEF *(RAC1^P29S^)*, WM1791 *(K-Ras^G12D^/RAC1^P29S^)* and WM1960 *(N-RAS^Q61K^/RAC1^P29S^)*. Table 2 displays the mutations of each melanoma cell line used in this study.

**Table 2.**
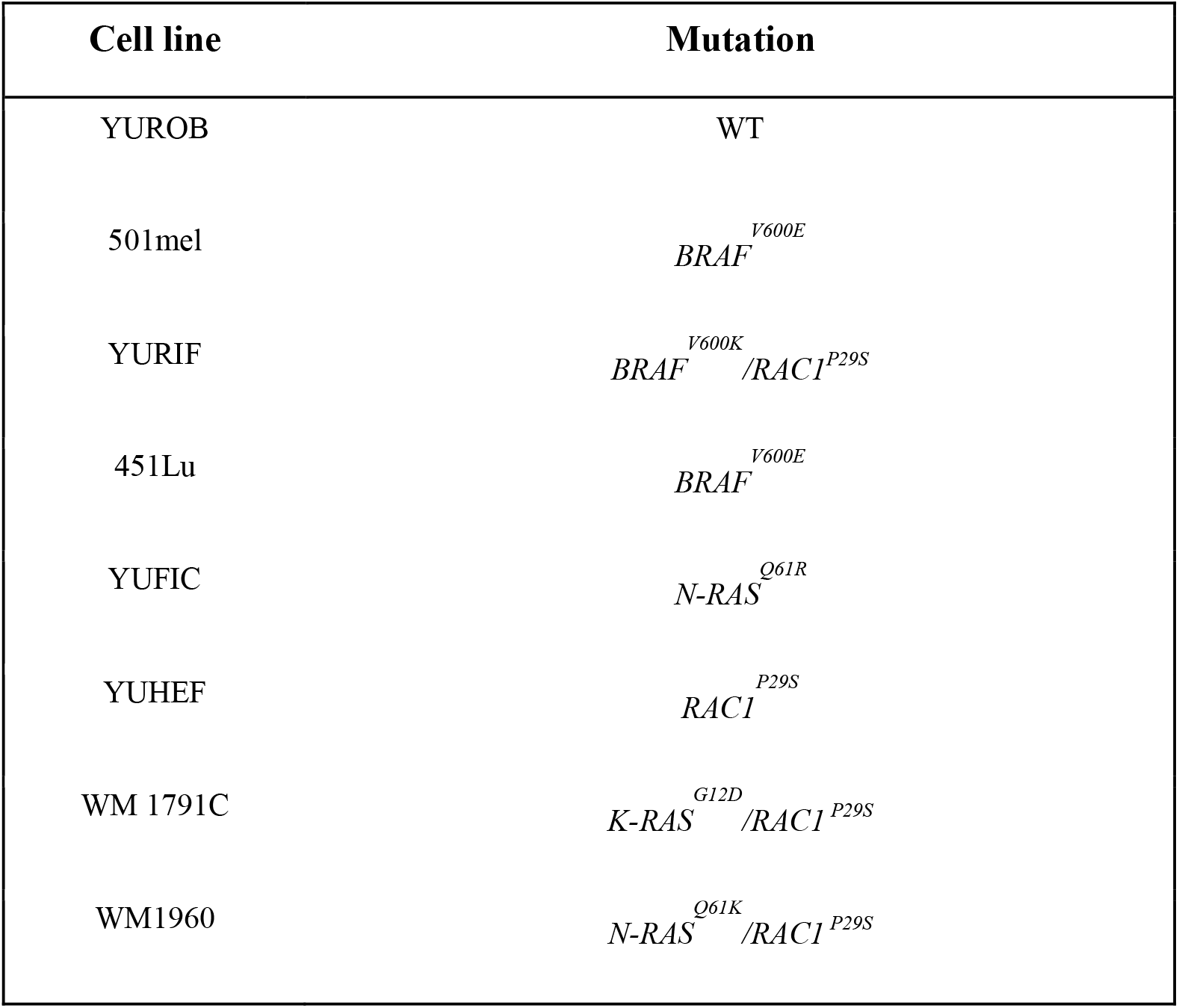
Genetic profiles of melanoma cell lines.

We evaluated the viability of melanoma cell lines when exposed to increasing concentrations of isoform-selective PI3K inhibitors as well a pan-PI3K inhibitor, using the reduction of MTT to formazan at 72 hours as readout. Viability was significantly reduced in all the melanoma cell lines exposed to the pan-PI3K inhibitor BKM120 (Fig. 1A). Using isoform-selective inhibitors, we found that PI3Kβ inhibitors (TGX221, GSK2636771) preferentially affected cell lines containing *RAC1* mutations (Fig. 1C and 1D) whereas the PI3Kα inhibitor (BYL719) decreased the viability of melanoma cells lacking this mutation (Fig. 1B). PI3Kγ and δ inhibitors (AS252424, GSK2269557) showed variable or no effects on melanoma cell lines, while affecting WT cell lines (Fig. 1E and 1F).

**Figure 1.**
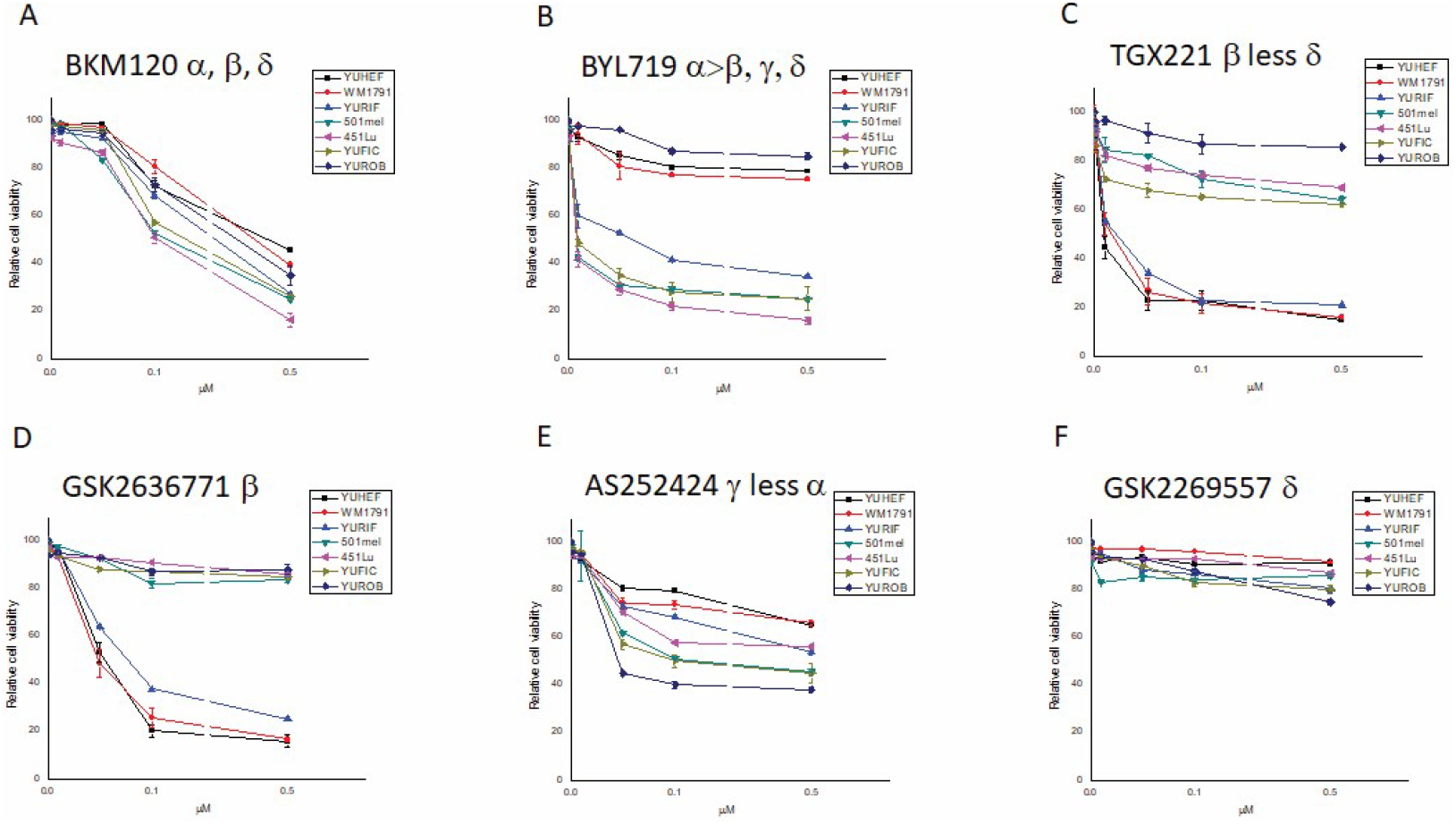
Effect of selective PI3K inhibitors on melanoma cell viability. Melanoma cells were treated for 72 h with concentrations ranging from 0.01 μM to 0.5 μM of A) pan-PI3K (BKM120), B) PI3Kα (BYL719), C) PI3Kβ (TGX221), D) PI3Kβ (GSK2636771), E) PI3Kγ (AS252424), F) PI3Kδ (GSK2269557). Cell viability was determined by an MTT assay.

### Cell proliferation and migration in response to PI3K inhibitors

We next evaluated the consequences of PI3K inhibition on cell proliferation and migration. Proliferation was studied in three melanoma cell lines with different genotypes: YUROB (WT), 501mel *(BRAF^V600E^)* and YUHEF *(RAC1^P29S^)*. WT cell proliferation was not affected by any of the PI3K or PAK1 (Frax-1036) inhibitors (Fig. 2). Treatment of RAC1^P29S^ mutant cell lines with the Group A PAK inhibitor Frax-1036 nearly abolished cell proliferation (Fig. 2, green line). PI3Kβ inhibitors (TGX221, GSK2636771) and pan-PI3K inhibitor (BKM120) showed an intermediate response, while the other drugs were less effective. In contrast, 501mel cells, bearing the BRAF^*V600E*^ mutation, displayed a reduced proliferation rate following treatment with pan-PI3K and PI3Kα inhibitors. Mild growth inhibition was observed when cells were exposed to the other PI3K inhibitors tested in this study.

**Figure 2.**
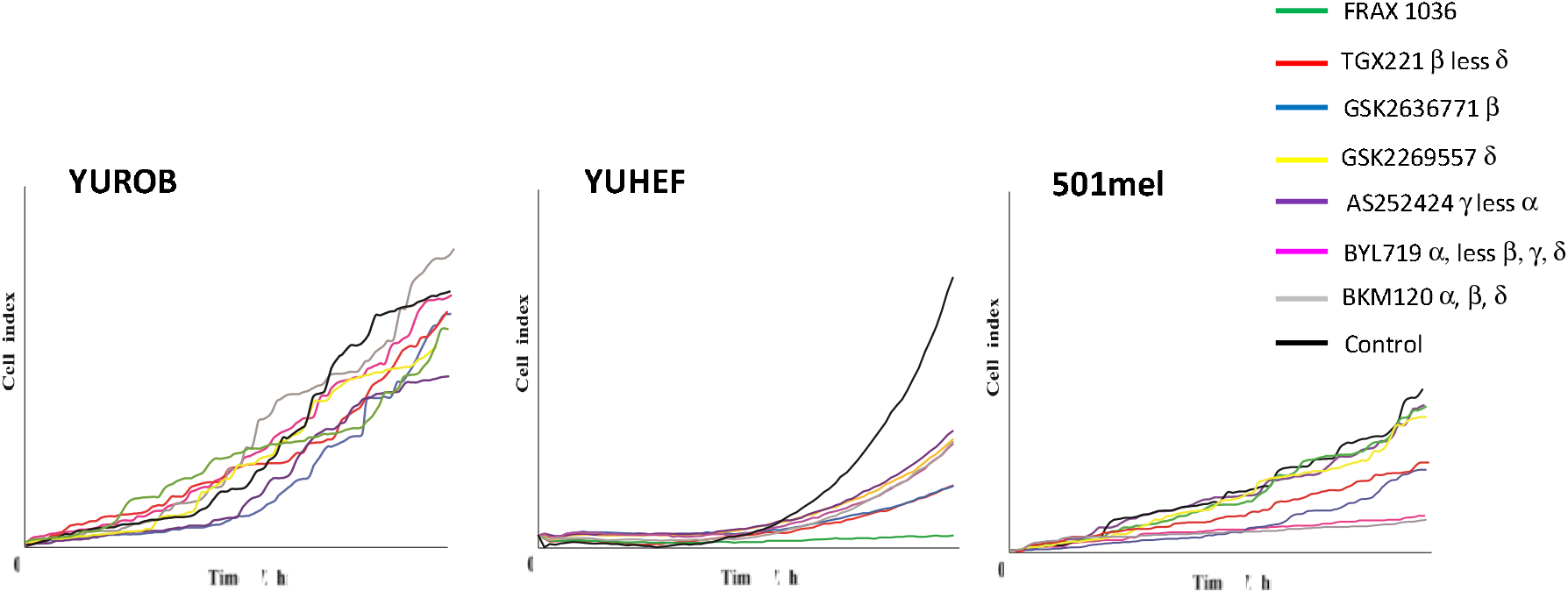
Proliferation of WT, *BRAF* and *RAC1*-mutant melanoma cell lines in presence of targeted inhibitors. Melanoma cells were treated with 100 nM of pan-PI3K (BKM120), PI3Kα (BYL719), PI3Kβ (TGX221, GSK2636771), PI3Kγ (AS252424), PI3Kδ (GSK2269557) and PAK1 (Frax-1036) small molecule inhibitors in A) WT cells (YUROB), B) RAC1 mutant (YUHEF) and C) BRAF mutant (501mel). Cell number was measured during 72h using an XCELLigence device.

RAC1 is well-known as a regulator of cell motility through its effects on the actin cytoskeleton. A wound-healing model was used to test whether PI3K inhibitors would differentially affect the migration of melanoma cells depending on their driver mutation(s). None of the inhibitors impeded migration in WT cell lines (YUROB). The migration of *RAC1* mutant cells (YUHEF) was severely diminished by the pan-PI3K and the PI3Kβ inhibitors, and partially reduced by the PI3Kα-inhibitor. The *BRAF* mutant cell (501mel) migration was reduced by exposure to the pan-PI3K and the PI3Kα inhibitors. When cell lines had both *RAC1* and *BRAF* mutations (YURIF), cells migration was strongly diminished by the pan-PI3K, the PI3Kα, and the PI3Kβ inhibitors. PI3Kγ and δ inhibitors were less potent in all the mutant cell lines (Fig. 3).

**Figure 3.**
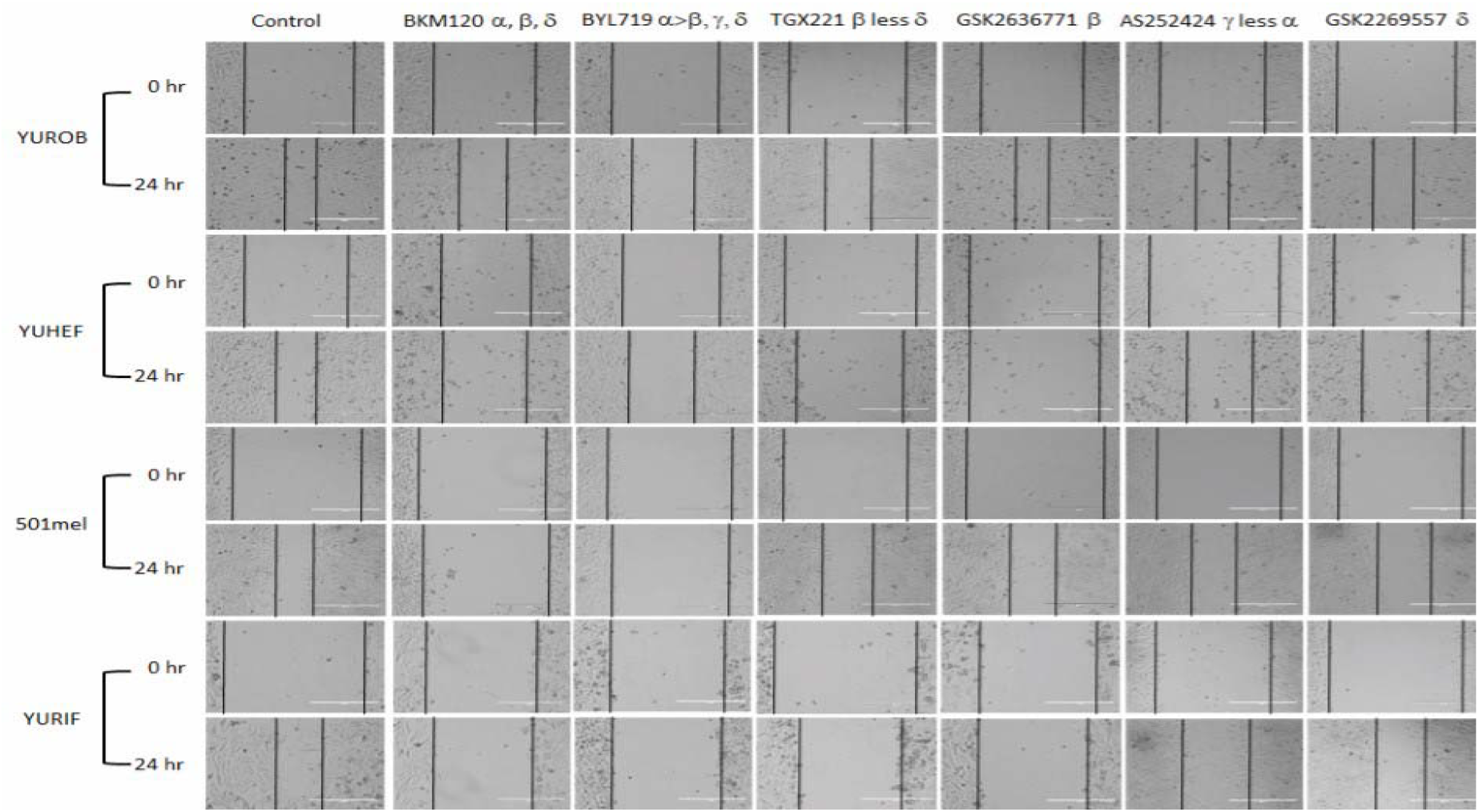
Consequences of PI3K inhibitors on cell migration. A cell culture wound closure assay was developed in six-well confluent plates. After the scratch, WT (YUROB), RAC1 (YUHEF), BRAF (501mel) and BRAF/RAC1 (YURIF) cell lines were treated with 100 nM of pan-PI3K (BKM120), PI3Kα (BYL719), PI3Kβ (TGX221, GSK2636771), PI3Kγ (AS252424) and PI3Kδ (GSK2269557) inhibitors. Snapshot images were taken in an inverted microscope at 0 and 24 h.

### RAC1-mutant signaling response to PI3K inhibitors

To assess the functional role of PI3K inhibition we examined RAC1 downstream effector proteins signaling in response to PI3K inhibitors in melanoma cell lines. Western blot analysis showed that treatment with specific PI3K drugs (α, β, γ, δ and pan-PI3K inhibitor) did not affect signaling in the WT cell line, but modified signaling in the mutant cell lines (Fig. 4). In non-treated cells, PAK and AKT activity was increased in *RAC1* mutants while ERK activity was constant in all the cell lines. The phosphorylation of PAK and AKT was significantly decreased in *RAC1* mutants by the pan-PI3K inhibitor (BKM120) and by the PI3Kβ inhibitors (TGX221 and GSK2636771). Additionally, the PI3Kα inhibitor (BYL719) significantly reduced the activation of AKT, as assessed by p-AKT blot. No effects were observed when cells where treated with the PI3Kγ and δ inhibitors. ERK activity was nearly abolished in all cell lines by the pan-PI3K inhibitor, and was suppressed by the PI3Kα inhibitor (BYL719) in BRAF-mutant cells and by PI3Kβ-inhibitors (TGX221 and GSK2636771) in RAC1-mutant cell lines. These results suggest that PI3K regulates signaling of AKT and PAK through different isoforms. (Fig. 4).

**Figure 4.**
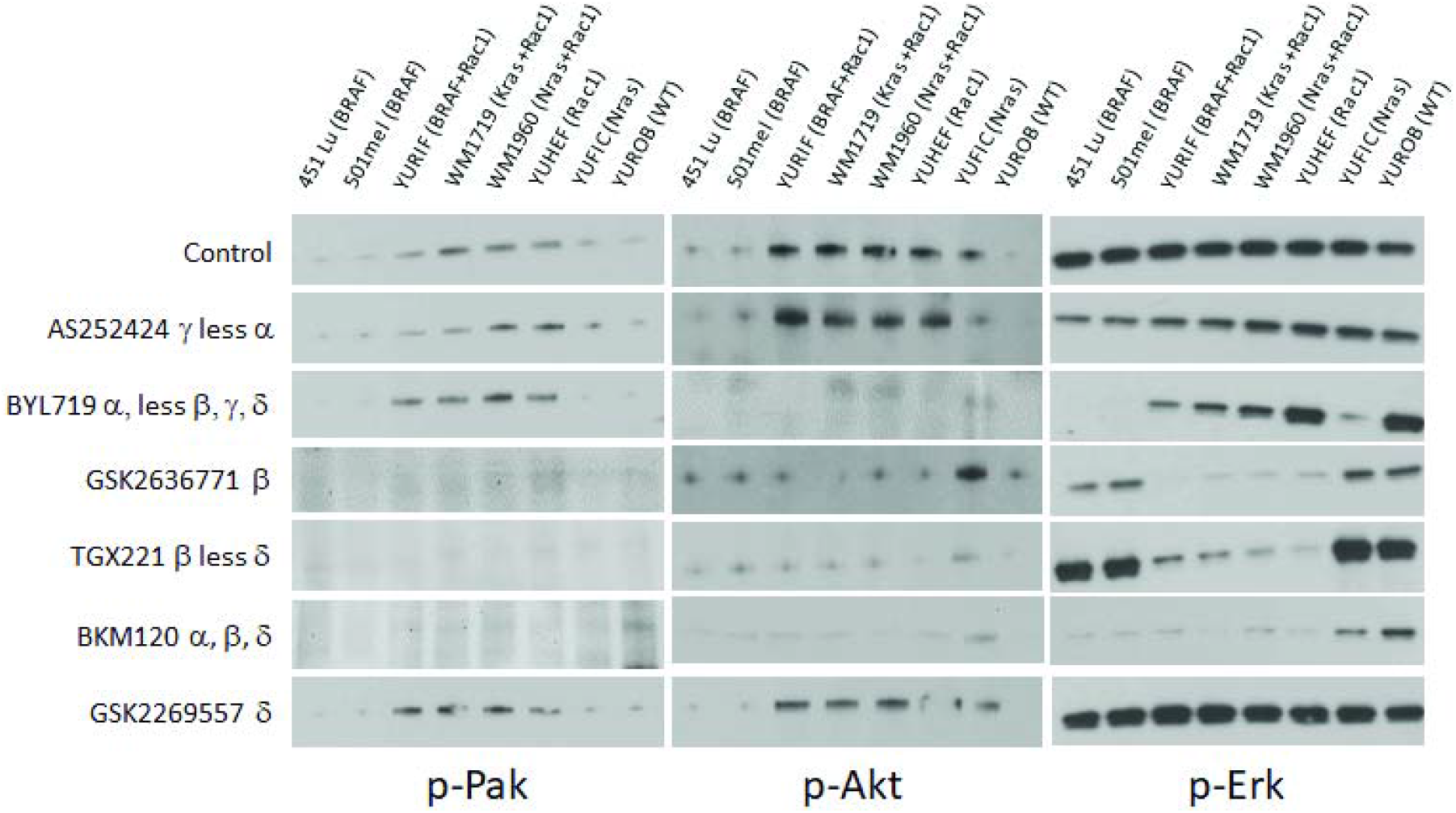
PI3K inhibitors cell signaling modification in melanoma cell lines bearing RAC1 and BRAF mutations. Melanoma cell lines with different genotypes were grown under standard conditions. Cells were treated with vehicle (control) or 100 nM of the indicated PI3K inhibitors for 24 h. Lysates were analyzed by Western blot for PAK, AKT, and ERK phosphorylation.

### Effect of inhibitors on RAC1 zebrafish embryonic development

To examine *in vivo* signaling roles for RAC1 effectors, we employed a zebrafish embryonic development assay. Overexpression of *RAC1^P29S^* hinders zebrafish embryonic development and activates ERK signaling^22^. We previously showed that these defects were blocked by inhibiting components of the RAS/MAPK pathway with small molecule PAK or MEK inhibitors^22^. Therefore, we asked whether these developmental effects could be suppressed by small molecule inhibitors of additional RAC1 effectors such as PI3K and SRF/MRTF. To determine if these inhibitors affect development, we introduced mRNA encoding *RAC1^P29S^* into one-cell zebrafish embryos. Abnormal RAC1 phenotype is characterized by pericardial edema, small/absent eyes, and reduced head size in ~97% of the embryos (Fig. 5). Specific small molecule inhibitors of PI3K, SRF/MRTF, AKT and PAK were diluted in the embryonic water at 4 hpf, washed out at 5 hpf, and development was followed for 24 hours. We demonstrated that a Group 1 PAK inhibitor (Frax-1036) (40/38 normal, 40/2 abnormal) and a MRTF/SRF inhibitor (CCG-203971) (40/33 normal, 40/7 abnormal) almost completely blocked the abnormalities induced by activated RAC1. In contrast, the PI3K inhibitors (TGX221, GSK2636771, BKM120 and BYL71) only partially prevented the Rasopathy-like phenotype; the pan-PI3K inhibitor, BKM120 (40/33 normal, 40/7 abnormal) and the PI3Kα, BYL71 (40/35 normal, 40/5 abnormal) had the best results. Finally, mutants treated with the AKT inhibitor had the least effect (40/24 normal, 40/16 abnormal) (Fig. 5). These results suggest that, of the three main groups of RAC1 effectors, Group A PAKS and/or SRF/MRTF represent the most effective targets to antagonize the developmental effects induced by RAC1^P29S^.

**Figure 5.**
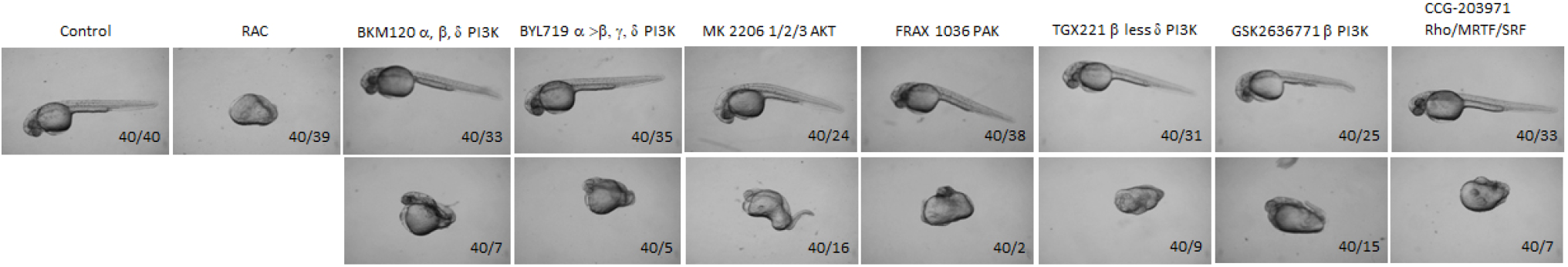
RAC1 overexpression in zebrafish embryonic development and its treatment with PI3K, AKT and Rho/MRTF/SRF inhibitors. One-cell zebrafish embryos were injected with RAC1^P29S^ mRNA. 1 μM inhibitors were added at 4 hpf, then removed and washed thoroughly. Embryonic morphology was scored at 24 hpf by a blinded observer. Quantitation of developmental abnormalities were performed with a Nikon digital sight DS Fi1 camera.

## Discussion

Melanoma is the most aggressive form of skin cancer and it is strongly associated with poor prognosis, low overall survival and drug resistance. RAC1^P29S^ is the third most common mutation found in sun-exposed melanoma. *RAC1* encodes a small GTPase that plays key roles in embryonic development, immune response, cell proliferation, survival, and rearrangement of cytoskeleton by actin filament remodeling.^3–5^ The Proline to Serine substitution in position 29 shifts the protein to a GTP-bound state activating its downstream effector protein PAK, PI3Kβ and SRF/MRTF among others. Furthermore, PI3Ks activate AKT, and have a positive feedback on RAC1 that may also activate some carcinogenic properties in cells.^14^

The inhibition of RAC1^P29S^ and its downstream effectors might provide new therapeutic targets for drug resistant tumors and advanced stages melanoma. Since few effective targets for RAC1 mutated melanoma, other than PAK1, have been reported, we examined the effect of PI3K and SRF/MRTF inhibition in melanocyte signaling and zebrafish development.

Cell viability and migration assays reveal that inhibiting each PI3K isoform yields different effects in NRAS, BRAF or RAC1 melanomas. The most effective inhibitors against RAC1 mutant cell viability were GSK2636771 and TGX221, which bind specifically to PI3Kβ. These inhibitors reduced RAC1^P29S^, KRAS^G12D^/RAC1^P29S^ and BRAF^V600K^/RAC1^P29S^ cell viability and are ineffective against WT, NRAS and BRAF melanoma cells. A PI3Kα inhibitor, BYL719, reduced viability of BRAF^V600K^/RAC1^P29S^, NRAS^Q61R^, and both BRAF^V600E^ cell lines and had no effect on WT, RAC1 and KRAS/RAC1 melanoma cells. As expected, the PI3Kδ and γ inhibitors had fewer effects on melanocytes cell viability since they are mainly expressed in leukocytes.^12^ This inhibitor-effect pattern is consistent in our proliferation and migration experiments. The pan-PI3K inhibitor showed a non-selective decline in cell viability, growth and migration.

We previously reported that the PAK inhibitor Frax-1036 blocks the growth and migration effects of an overactive RAC1 GTPase in cell lines; and reverse the rasopathy-like phenotype shown in zebrafish when injected with RAC1 mRNA^9^. As expected, the PAK inhibitor effectively prevented the developmental effects caused by RAC1^P29S^. PI3K inhibitors were in general less effective, with some isoform specificity: 17.5% of the embryos incubated with the pan-PI3K presented dysfunctional development; 12.5% when using the PI3Kα inhibitor; 22.5% and 37.5% when using the PI3Kβ inhibitors TGX221 and GSK2636771 respectively. When treated with the SRF/MRTF inhibitor, only 17% of the embryos presented severe development impairments. The AKT inhibitor was the least efficient agent to inhibit the developmental effects of RAC1^P29S^.

While the pan-PI3K inhibitors impede growth and migration in all melanoma cell lines tested, PI3Kβ small molecule inhibitors specifically impede RAC1-, while PI3Kα specifically impedes BRAF-driven melanoma cell line proliferation. In embryonic zebrafish development, the rasopathy-like phenotype induced by mutant RAC1 was most successfully reversed when using the PAK or the SRF/MRTF inhibitor, and to a lesser degree by the pan-PI3K, the PI3Kα, and the PI3Kβ inhibitors (TGX221, see Table 1). These results, like those recently reported in a *Rac1^P29S^* mouse model^11^, suggest that targeting Group A PAKs and/or SRF/MRTF, and possibly also PI3K/AKT signaling, could become a promising approach to suppress RAC1 signaling in malignant melanoma.

## Disclosure of Potential Conflicts of Interest

No potential conflicts of interest were disclosed.

## Funding

This work was supported by grants from the NIH (JC), the Melanoma Research Foundation (JC), and Presupuesto Interno del Instituto de Química/UNAM (DAO) as well as Cátedras CONACYT and NCI Core Grant P30 CA06927 (to Fox Chase Cancer Center, FCCC).

## Acknowledgments

We thank Drs. Ruth Halaban and Meenhard Herlyn for providing melanoma cell lines, Genentech for providing Frax-1036, and the Fox Chase Cancer Center Animal Facility for assistance with zebrafish experiments.

## Declaration of Interests

The authors claim no conflicts of interest.

